# Dual blockade of IL-6 and CTLA-4 regresses pancreatic tumors in a CD4^+^ T cell-dependent manner

**DOI:** 10.1101/2020.02.07.939199

**Authors:** Michael Brandon Ware, Christopher McQuinn, Mohammad Y. Zaidi, Hannah Knochelmann, Thomas A. Mace, Zhengjia Chen, Chao Zhang, Matthew R. Farren, Amanda N. Ruggieri, Jacob Bowers, Reena Shakya, A. Brad Farris, Gregory Young, William E. Carson, Bassel El-Rayes, Chrystal M. Paulos, Gregory B. Lesinski

**Affiliations:** Department of Hematology and Medical Oncology, Winship Cancer Institute of Emory University; Division of Surgical Oncology, Department of Surgery, Department of Internal Medicine, The Ohio State University; Department of Surgery, Winship Cancer Institute of Emory University; Department of Microbiology and Immunology, Hollings Cancer Center, Medical University of South Carolina; Division of Gastroenterology Hepatology and Nutrition, Department of Internal Medicine, The Ohio State University; Department of Biostatistics, Emory University; Comprehensive Cancer Center, The Ohio State University; Department of Pathology, Winship Cancer Institute of Emory University; Center for Biostatistics, The Ohio State University

**Author notes:** To whom correspondence should be addressed: Gregory B. Lesinski, Department of Hematology and Medical Oncology, Winship Cancer Institute of Emory University, 1365 Clifton Rd. NE, Atlanta, GA, 30322, USA.

## Abstract

Pancreatic ductal adenocarcinoma (PDAC) is exceptionally resistant to immune checkpoint inhibition (ICI). We previously reported that elevated systemic interleukin-6 (IL-6) and increased numbers of T cells positive for circulating cytotoxic T-lymphocyte-associated protein 4 (CTLA-4) correlate with worse overall survival in patients with PDAC. We postulated that combined blockade of IL-6 and CTLA-4 would significantly enhance anti-tumor immune responses to PDAC. Dual blockade of IL-6 and CTLA-4 in immune competent mice bearing subcutaneously injected pancreatic tumors significantly inhibited tumor growth, accompanied by overwhelming T cell infiltration. Therapeutic efficacy was confirmed in an orthotopic murine model of pancreatic cancer and T cell depletion studies unveiled a unique dependence on CD4^+^ T cells for anti-tumor activity of dual IL-6 and CTLA-4 blockade. *In vitro* studies utilizing T cells from a TRP-1 transgenic mouse as an antigen-specific model system demonstrate this combination therapy elicits increased IFN-γ production by activated CD4^+^ T cells. Additionally, IFN-γ stimulation of pancreatic tumor cells *in vitro* profoundly increased tumor cell production of CXCR3 specific chemokines (CXCL10 and CXCL9). Further studies blocking CXCR3 in the presence of combined IL-6 and CTLA-4 blockade prevented orthotopic tumor regression, demonstrating a dependence on the CXCR3 axis for anti-tumor efficacy. We also found combination therapy increased intratumoral CD4^+^ T cells and elicited systemic changes in T-helper subsets. These data represent the first report of IL-6 and CTLA-4 blockade as a means to regress pancreatic tumors with defined operative mechanisms of efficacy. Given these results, this therapeutic combination has potential for immediate clinical translation.

**One Sentence Summary:** Blockade of interleukin-6 in pancreatic cancer enhances CTLA-4 immune checkpoint inhibition to regress tumors in a CD4^+^ T cell and CXCR3-dependent manner.

## Introduction

Antibodies targeting immune checkpoint receptors and their ligands have gained regulatory approval and demonstrated clinical efficacy in patients with a number of malignancies. The most notable examples include blockade of cytotoxic T-lymphocyte protein-4 (CTLA-4) and programmed cell death protein 1 (PD-1) or its ligand, programed cell death protein ligand 1 (PD-L1). Despite encouraging data in patients with various malignancies, including melanoma, bladder, non-small cell lung, and microsatellite instable (MSI) cancers among others, there remain a number of key challenges with this approach (*1*). For example, many patients still do not gain clinical benefit from immune checkpoint inhibition (ICI), while resistance is common in those who do initially respond to therapy (*1*). Most tumors arising in the pancreas and other portions of the gastrointestinal system, are also inherently resistant to ICI (*2*). Overcoming the limitations for individuals unresponsive to these emerging therapies is a priority area of research, as it could advance treatment outcomes across multiple tumor types.

Combination approaches involving immune checkpoint blockade are quickly expanding with promising results. Data are emerging from early phase clinical trials combining immune checkpoint blockade with conventional cytotoxic or radiation therapy, concurrent modulation of multiple immune checkpoint receptors or ligands, or together with targeted small molecules (*3–7*). The intent of these approaches is to overcome redundant immune suppressive mechanisms that limit T cell mediated immunity or concurrently act upon the malignant cells. Given the number of ongoing studies, it is likely that novel strategies to overcome resistance to immunotherapy will continue to materialize. Research into the mechanisms of resistance may lead to the application of tailored therapeutic strategies based on the underlying immune suppressive mechanisms.

The highly desmoplastic stroma unique to PDAC is a dynamic immune suppressive component contributing to the poor impact of immune therapy in this malignancy. Our laboratory and others have recently demonstrated that the pancreatic cancer stroma and stromal-derived cytokines restrain host immunity (*8–11*). Although dysregulated cytokines represent rational therapeutic targets, there are limited data to help prioritize them in patients for immediate translation. Recent clinical studies support interfering with IL-6 as a therapeutic target to enhance immune checkpoint blockade in patients with advanced malignancy (*12*). In a cohort of seventy-two treatment naïve patients with metastatic pancreatic ductal adenocarcinoma, circulating IL-6 independently correlated with reduced overall survival (*13*). These data were intriguing as IL-6 can regulate phenotypic and functional properties of a smattering of various lymphocyte and myeloid cell populations (*14, 15*). Detailed immune phenotyping of peripheral blood mononuclear cells from the same cohort of patients revealed additional observations of interest. Notably, a similar relationship between reduced overall survival and elevated circulating T cells expressing CTLA-4 was observed (*13*). These data encourage strategic combination therapies incorporating CTLA-4 targeting antibodies in PDAC

Extensive mechanistic studies have revealed that inhibition of the cytokine, interleukin-6 (IL-6), derived from pancreatic stellate cells (PSC) in the TME, enhanced PD-L1 blockade therapy (*16, 17*). In fact, dual blockade of IL-6/IL-6 receptor signaling and PD-L1 led to CD8^+^ T cell dependent anti-tumor immunity. This combination therapy increased the infiltration of effector T cells into tumors, and systemically expanded cells with a Th1 T cell phenotype. These data suggest that targeting tumor or stromal-derived cytokines may be an additional way to enhance the efficacy of immunotherapy in patients with otherwise refractory malignancies. To our knowledge, concurrent targeting of immune suppressive cytokines and immune checkpoints represents an area with limited experimental investigation, yet has high potential for rapid translation into patients. Whether or not IL-6 blockade will augment efficacy of immune checkpoint inhibitors other than PD-1/PD-L1 remains an important unanswered question. Similarly, the mechanism(s) of action may differ depending on what targeting approaches this cytokine-blocking strategy is paired with.

Based on these data, we hypothesized that IL-6 blockade would enhance the efficacy of CTLA-4 blockade therapy. Herein, we report that combined blockade of IL-6 and CTLA-4 inhibits pancreatic tumor growth by potentiating the infiltration of T cells into tumors. Surprisingly, this therapeutic strategy is reliant on CD4^+^ T cell subsets for its efficacy. *In vitro,* this therapy promotes IFN-γ production by activated antigen-specific CD4^+^ T cells. In turn, we demonstrate IFN-γ promotes the production of lymphocyte-attracting chemokines by tumor cells, including high levels of the CXCR3-associated chemokine CXCL10. Further, this treatment regimen induced systemic shifts in CD4^+^ T-helper subsets and was dependent upon CXCR3 for efficacy. Together our new findings suggest that combined blockade of IL-6 and CTLA-4 can regress pancreatic tumors via a unique mechanism by imparting CD4^+^ T cell-mediated anti-tumor immune responses.

## Results

### Combined blockade of IL-6 and CTLA-4 augments antitumor efficacy in murine pancreatic tumor models

Increased levels of the pleiotropic cytokine IL-6 and higher numbers of CTLA-4^+^ T cells are directly linked to reduced survival in patients with metastatic PDAC (*13*). Thus, we rationalized that targeting IL-6 may enhance the efficacy of ICI. We first sought to determine the efficacy of dual IL-6 and CTLA-4 blockade in mice bearing subcutaneous MT5 tumors. The MT5 cell line is relevant to human patients, as it originated from KPC tumors that harbor both G12D mutated *Kras* and R172H mutated *Trp53* (*18*). Tumor growth was significantly reduced in mice treated with combined IL-6 and CTLA-4 blocking antibodies. As expected, mice treated with isotype control antibodies rapidly succumb to disease compared to those given the dual therapy (p=0.0001) (**Figure 1A**). Although single agent anti-CTLA-4 inhibited tumor growth to a greater extent than mice treated with an isotype control (p<0.0004), consistent with our prior studies (*17*), blockade of IL-6 alone did not delay tumor growth. Notably, mice receiving the combination of antibodies to IL-6 and CTLA-4 together experienced more impressive tumor growth inhibition compared to mice receiving single agent antibodies to CTLA-4 (p=0.0207) or IL-6 (p=0.0002). This dramatic impact on tumor growth was encouraging, however the mechanism of action leading to this result was unknown.

**Figure 1.**
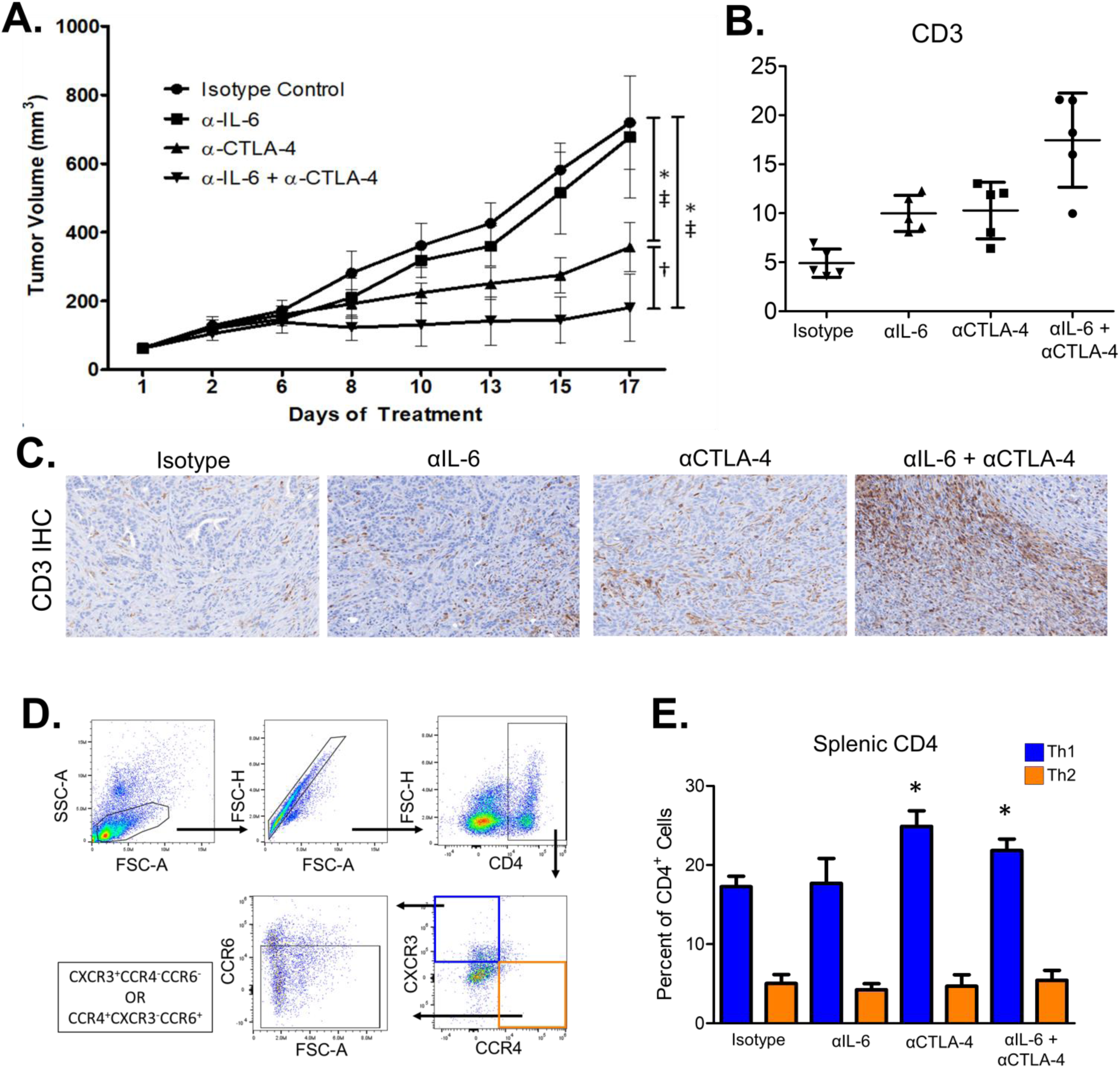
Combined blockade of IL-6 and CTLA-4 significantly inhibits tumor growth and promotes CD8 T cell infiltration of tumors in a subcutaneous murine model of pancreatic cancer. MT5 murine pancreatic tumor cells were subcutaneously injected into C57BL/6 mice with treatment beginning when tumors reached 50-100mm^3^. Mice were treated with 200mg (intraperitoneal injection 3 times/week) of isotype control, cytokine blockade (anti-IL-6) and/or anti-CTLA-4 antibodies (n=5 mice/group) until mice met pre-specified IACUC-approved early removal criteria. **(A)** Changes in tumor volume as determined by caliper measurement throughout the course of antibody treatment. Mean ± SD; * P<0.05 vs. Isotype Control, ‡ P<0.05 vs. Anti-IL-6, † P<0.05 vs. Anti-CTLA-4. **(B)** Representative 20x images of IHC staining for CD3 in FFPE tumor tissue slices from mice in the different treatment groups. **(C)** Mean ± SD for percent of cells expressing CD3^+^ in subcutaneous tumors per high-powered field. Symbols represent individual mice; * indicates significance compared to isotype control treated mice. **(D)** Splenocytes were isolated from the mice receiving treatment as stated in figure 1A. Flow cytometry was performed with antibodies against CD4, CCR6, CXCR3, CCR4, and RORγt. CD4^+^CCR6^-^CXCR3^+^CCR4^-^ were identified as suggestive of a Th1 phenotype and CD4^+^CCR6^-^ CXCR3^-^CCR4^+^ as a Th2 phenotype. **(E)** Graph of mean percentages of CD4^+^ T cells that have a Th1 or Th2 phenotype. Data shown as Mean ± SD; * indicates significance compared to isotype control treated mice.

### Combined blockade of IL-6 and CTLA-4 mediates increased T cells in pancreatic tumors

In light of previous research by ourselves and others investigating the influence of IL-6 and CTLA-4 on T cell populations (*17, 19*), we hypothesized this combined blockade could impact T cell infiltration into pancreatic tumors. Immunohistochemical staining of tumors from mice treated with this therapy indicated an altered presence of T cells in the tumor microenvironment (representative staining shown in **Figure 1B)**. Further quantification revealed that both single agent blockade of IL-6 (p=0.0038) or CTLA-4 (p=0.0035) increased the infiltration of CD3^+^ T cells as compared to tumors from isotype control-treated mice (**Figure 1C**). Mice given combined therapy had more T cells infiltrating the tumor compared to those mice treated with either single agent blockade of IL-6 (p=0.035), CTLA-4 (p=0.038) or isotype controls (p<0.0001) **(Figure 1C).**

### Systemic changes in Th1 immunity occur following combined IL-6 and CTLA-4 blockade

IL-6 plays a role in regulating differentiation and activation of T cell subsets (*20–22*). Therefore, we evaluated the impact of this combination treatment regimen on splenic-derived T cells as a surrogate of systemic immune response. Flow cytometric analysis of splenocytes obtained at the endpoint of the study described above revealed a number of phenotypic alterations that may influence anti-tumor immune responses. First, treatment with antibodies targeting CTLA-4 alone or in combination with IL-6 blockade increased circulating cells with a Th1 phenotype (CD4^+^CCR6^-^CXCR3^+^CCR4^-^) as compared to mice treated with isotype control (p<0.05) or antibodies targeting IL-6 alone (p<0.05) (**Figure 1D-E).** No change was observed in cells with a Th2 (CD4^+^CCR6^-^CXCR3^-^CCR4^+^) or Th17 (CD4^+^RORγt^+^) phenotype **(Figure 1D-E and Supplementary Figure 1A-B respectively).** Unexpectedly, the proportion of splenocytes expressing T-regulatory cell (Treg) phenotypic markers (CD4^+^CD25^+^FoxP3^+^) was higher in the combination treatment group in comparison to both isotype control (p<0.05) and anti-IL-6 (p<0.05) (**Supplemental Figure 1C-D**). Because IL-6 also regulates the expansion of myeloid-derived suppressor cells in pancreatic cancer (*16*), phenotypic properties of these cells were assessed. No significant changes in the frequency of either monocytic (CD11b^+^Ly6G^-^Ly6C^+^) or granulocytic (CD11b^+^Ly6G^+^Ly6C^low^) populations were observed across the individual treatment groups (**Supplemental Figure 1E-F**). These data indicate a dual role for combined blockade of IL-6 and CTLA-4 in driving T cell infiltration of tumors, while simultaneously altering T cell phenotypes. Given the influence of this therapy on Th1 immunity, we sought to determine if there might be any mechanistic connection between T cell infiltration and changes in immune phenotypes observed in mice receiving this combination therapy.

### CD4^+^ and CD8^+^ cells contribute to the efficacy of combined IL-6 and CTLA-4 blockade

Given the increased infiltration of CD3^+^ T cells into subcutaneous tumors and systemic alteration of CD4^+^ T cell subsets, we next questioned whether this combination therapy might be CD4^+^ or CD8^+^ T cell dependent. For these studies, we employed a more clinically-relevant model of murine pancreatic cancer by orthotopically implanting luciferase-expressing KPC-luc cancer cells into the pancreas of immune competent mice. These cells express an enhanced firefly luciferase construct that allows for longitudinal bioluminescent imaging (BLI) of tumors. Antibody depletion studies were conducted to investigate the individual role of CD4^+^ and CD8^+^ T cells in mediating the efficacy of combined IL-6 and CTLA-4 blockade. For these experiments, CD4^+^ or CD8^+^ T cells were depleted in mice bearing orthotopic KPC-luc tumors prior to treatment with isotype control antibodies, or combined IL-6 and CTLA-4 blockade (**Figure 2A**). Confirmation of CD4 or CD8 depletion was accomplished by flow cytometric analysis of CD4^+^ and CD8^+^ T cells within CD3^+^ T cells isolated from the spleens of mice at the study endpoint (**Figure 2B**).

**Figure 2.**
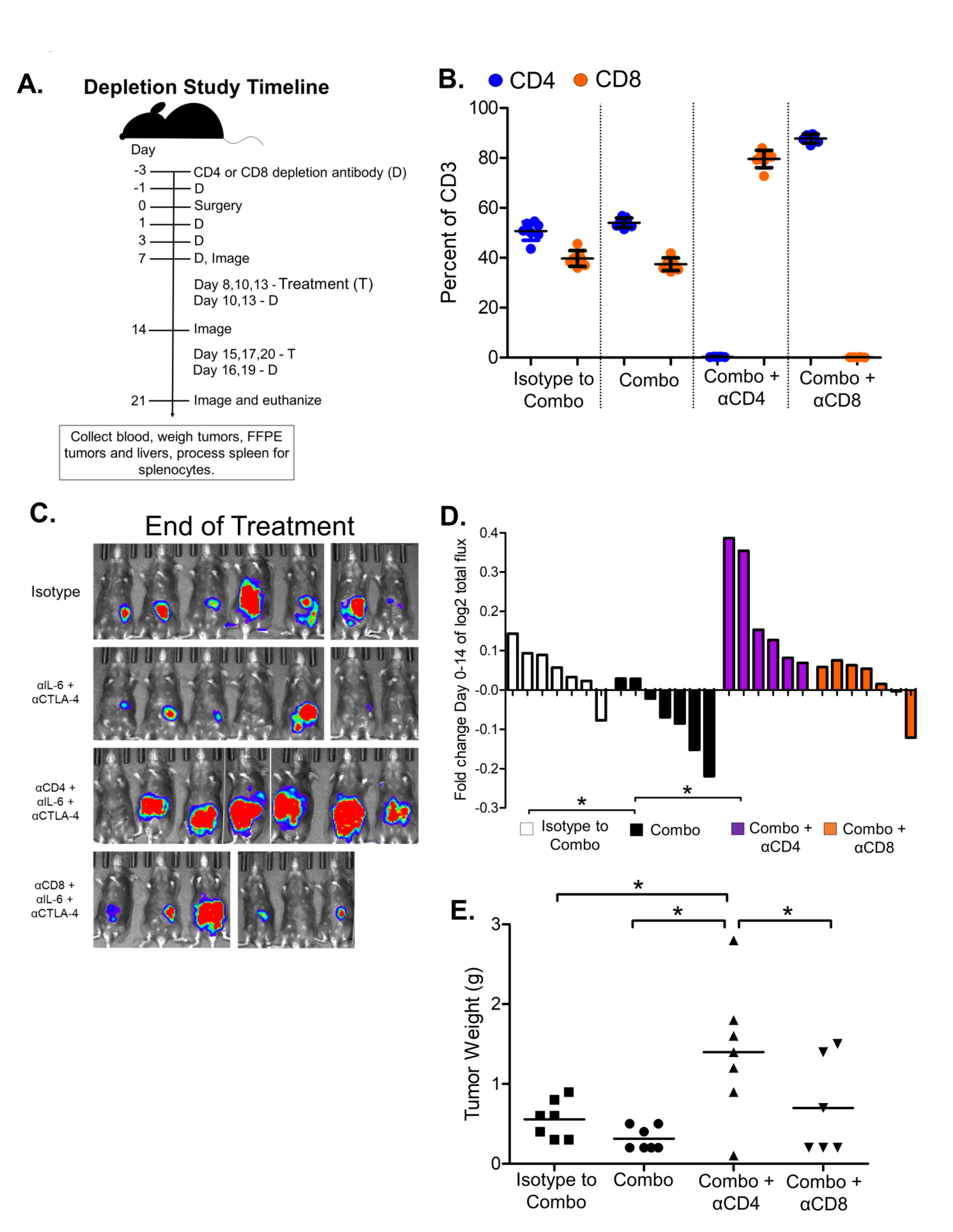
Both CD4^+^ and CD8^+^ T cells are required for anti-tumor responses to orthotopic pancreatic tumors in mice treated with combined IL-6 and CTLA-4 blockade. Female C57BL/6 mice, 6-8 weeks of age were orthotopically injected with 2×10^5^ KPC-luc cancer cells and imaged 1 week later by bioluminescent imaging (BLI) to confirm tumor establishment. **(A)** Timeline for the administration of CD4 or CD8 depleting antibodies relative to orthotopic injection and subsequent administration of IL-6 and CTLA-4 blocking antibodies or isotype control antibodies. **(B)** At the study endpoint, mice were euthanized and spleens were collected and processed for isolation of splenocytes. Splenocytes were then stained for CD3, CD4 and CD8 markers. The graph demonstrates the percentage of CD3^+^ cells in splenocytes from each group that expressed the markers CD4 (blue) or CD8 (orange). **(C)** EOT Bioluminescent images for the each mouse in the study outlined in Figure 2A are displayed. **(D)** Tumor growth for each mouse from the study outline in Figure 2A was measured over time by BLI and the fold change in Log^2^ of total flux for each mouse was graphed as a bar. **(E)** At the study endpoint (see outline in Figure 2A), mice were euthanized and the weight of each tumor was measured and graphed with symbols representing individual mice and mean displayed for each treatment group.

Longitudinal BLI data indicated the efficacy of combined IL-6 and CTLA-4 blockade to be CD4^+^ T cell-dependent, as mice depleted of CD4^+^ T cells receiving these therapeutic antibodies had accelerated tumor progression compared to mice receiving the combination therapy (**Figure 2C-D**). Tumor progression in some animals was striking, and faster than that of mice receiving isotype control antibodies. CD8^+^ T cell depletion also impacted tumor growth, albeit not to the magnitude of CD4^+^ T cell depletion, nor to a significant degree compared to combination treated mice (**Figure 2C-D**). To complement the trends observed with BLI data, the total pancreas and tumor weight was also measured post-mortem at the study endpoint (**Figure 2E**). These data confirmed consistent efficacy of the therapeutic combination and highlighted a unique coupled mechanism of action requiring CD4^+^ T cells.

### Combined IL-6 and CTLA-4 blockade supports Th1 cytokines that cross-talk to facilitate chemokine production from tumor cells

We posited that combined blockade of IL-6 and CTLA-4 would promote the generation of CD4^+^ T cells with a Th1 cytokine profile. To test this idea, we assayed how modulation of IL-6 and CTLA-4 impacted the biology of transgenic TRP-1 CD4^+^ T cells bearing a TCR that recognizes tyrosinase-related protein (TRP-1) as a means to model recognition of endogenous tumor antigen. Consistent with our findings in the MT5 pancreatic model, we found that expanding TRP-1 CD4^+^ T cells in the presence of IL-6 and CTLA-4 blocking antibodies fostered their capacity to secrete IFN-γ when re-stimulated with cognate antigen (**Figure 3A-B)**. This work suggests that dual blockade therapy imparts a direct effect on T cells, which in turn drives production of Th1 associated cytokines by CD4^+^ T cells.

**Figure 3.**
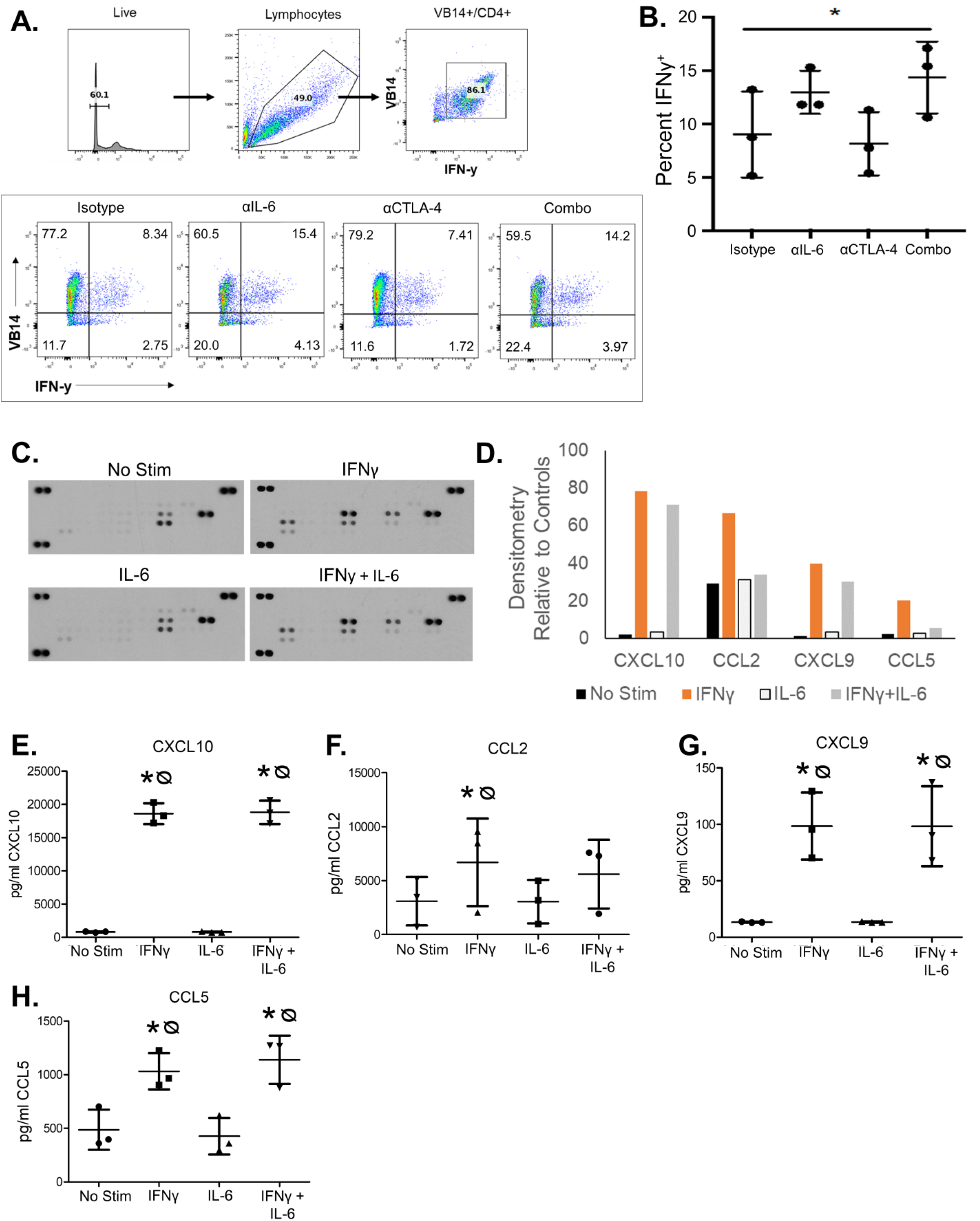
Combined blockade of IL-6 and CTLA-4 promotes IFN-γ production by antigen-activated CD4 T cells, which can elicit changes in chemokine production by pancreatic tumor cells. **(A)** Trp-1 specific CD4^+^ T cells were stimulated in the presence of antibodies to IL-6, CTLA-4, the combination of both or isotype control antibodies. The percentage of cells expressing Vβ14 and IFN-γ were quantified by flow cytometry. **(B)** Graph shows percentages of CD4^+^ T cells expressing IFN-γ for each mouse with mean ±SD. CD4^+^ T cells expressed more IFN-γ in the presence of dual CTLA-4 and IL-6 blockade compared to isotype control antibodies (p=0.0217) **(C)** KPC-luc cells were plated at 2×10^5^ cells per well in 6 well plates and then stimulated with 1µg/ml IFNy, 10ng/ml IL-6, both or vehicle control for 24 hrs. Resulting supernatants were collected and analyzed using the Proteome Profiler Mouse Chemokine Array Kit. Shown are resulting images of a chemokine membrane exposed to supernatants of KPC-luc cells from each treatment condition. **(D)** The relative densitometry to loading controls for CXCL10, CCL2, CXCL9, CCL5 as detected by the chemokine array are graphed for each treatment condition. Repetitions of the experiment described in Figure 3C were quantified by ELISA and the resulting concentrations were graphed for **(E)** CXCL10 **(F)** CCL2 **(G)** CXCL9 and **(H)** CCL5. Data are shown with symbols marking individual experiments and mean ± SD for each group.

Next, we hypothesized that abundant Th1 cytokines induced by our dual therapy may modulate tumor-derived production of chemokines to attract T cells. To test this mechanism, we surveyed chemokine production from murine MT5 or KPC-Luc tumor cells (harboring mutated *Kras* and *tp53*) following *in vitro* stimulation with IFNγ, IL-6 or both cytokines together. Stimulation with IFN-γ resulted in canonical upregulation of chemokines from tumor cells, as detected by a chemokine array (**Supplemental Figure 2A**) including elevated CXCL10 and CXCL9, which ligate the CXCR3 receptor (**Figure 3C-D)** (*23–26*). CCL2 and CCL5 were also upregulated upon IFN-γ stimulation (**Figure 3C-D**). When tumor cells were concurrently stimulated with IL-6 and IFN-γ, we found that CCL2 and CCL5 production by tumor cells was dramatically decreased (**Figure 3C-D**). This data suggests that elevated IL-6 may interfere with Th1-cytokine mediated chemokine production by tumor cells. We repeated these experiments with KPC-luc (**Figure 3E-** and MT5 cells (**Supplemental Figure 2B-E**) and ELISA confirmed significant upregulation of CXCL10, CXCL9, CCL2 and CCL5 by treatment with IFN-γ. However, treatment of cells with IL-6 alone did not change chemokine production, nor did combined stimulation with IL-6 and IFN-γ differ significantly from IFN-γ stimulation alone as quantified by ELISA (**Figure3 E-H**).

### CXCR3 is required for the efficacy of combined IL-6 and CTLA-4 blockade

We hypothesized that improved Th1 cell trafficking into the PDAC tumor microenvironment via CXCR3 is a key mechanism that contributes to the efficacy of combined IL-6 and CTLA-4 blockade. To determine if CXCR3 receptor interactions were essential mediators of the observed T cell response, we blocked CXCR3 via administration of blocking antibodies in mice receiving our combination therapy. No significant change in tumor growth rate was detected by BLI when comparing isotype control mice to those receiving single agent therapy. However, all mice receiving combined blockade of IL-6 and CTLA-4 experienced tumor regression as detected by BLI (**Figure 4A-B**). This change in BLI signal of mice receiving combination therapy was significant compared to mice receiving isotype control antibodies, or single agent blockade of IL-6 or CTLA-4. Concurrent CXCR3 blockade significantly inhibited the efficacy of the combination therapy, leading to results similar to that seen when mice were treated with isotype controls (**Figure 4A-B**). Analysis of post-mortem pancreas/tumor weight at the study endpoint confirmed growth inhibitory effects of the combination therapy were significant, as compared to treatment with either CTLA-4 blockade or isotype control antibodies (p<0.05; **Figure 4C**). Histologic analysis was utilized to survey changes in the tumor microenvironment that may explain the dramatic efficacy of this treatment combination. Although prior observations from our group show IL-6 is largely derived from fibroblasts in the PDAC tumor microenvironment (*16, 17*), no difference in alpha smooth muscle actin (α-SMA) staining was observed in tumors between treatment groups (**Supplemental Figure 3A-B**). A trend toward increased CD8^+^ T cells in tumors from mice receiving dual blockade was seen, but did not reach significance (**Figure 5A-B**). Although no differences in CD4^+^FoxP3^+^ T cells emerged (**Supplemental Figure 3C**), a profound increase in CD4^+^ T cells lacking FOXP3 expression was observed in tumors from mice receiving combined IL-6 and CTLA-4 blockade compared to control mice (p=0.0297) or to mice receiving IL-6 single agent blockade (p=0.0439) (**Figure 5C-D**).

**Figure 4.**
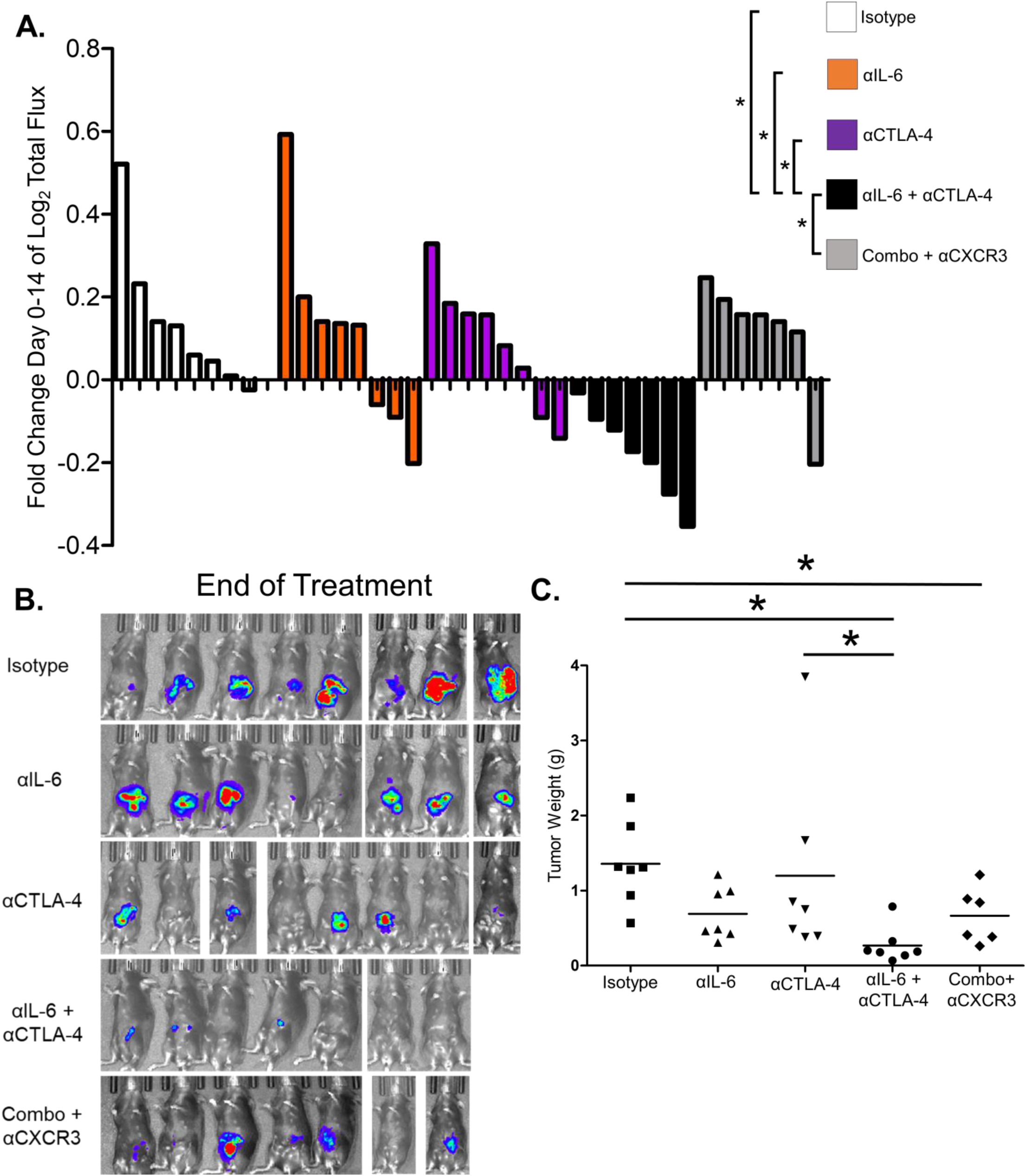
Treatment of murine orthotopic pancreatic tumors with antibodies to IL-6 and CTLA-4 results in significant tumor regression and increased intra-tumoral CD4^+^ and CD8^+^ T cells in a CXCR3 dependent manner. Female C57BL/6 mice, 6-8 weeks of age were orthotopically injected with 2×10^5^ KPC-luc cancer cells and imaged 1 week later by bioluminescent imaging (BLI) to confirm tumor establishment. Mice were then treated 3 times a week for 2 weeks with antibodies to IL-6, CTLA-4, the combination of both, the combination and antibodies to CXCR3 or isotype control antibodies. **(A)** Tumor growth was measured over time by BLI and the Log^2^ fold change in total flux for each mouse was graphed as a bar. * indicates significance (p<0.05) to mice receiving dual blockade of IL-6 and CTLA-4. **(B)** Resulting bioluminescent images for each mouse at the end of treatment demonstrates the anti-tumor efficacy measured by BLI. **(C)** After two weeks of treatment, mice were euthanized and the total weight of each tumor was collected and graphed as mean for each treatment group. Symbols represent individual mice.

**Figure 5.**
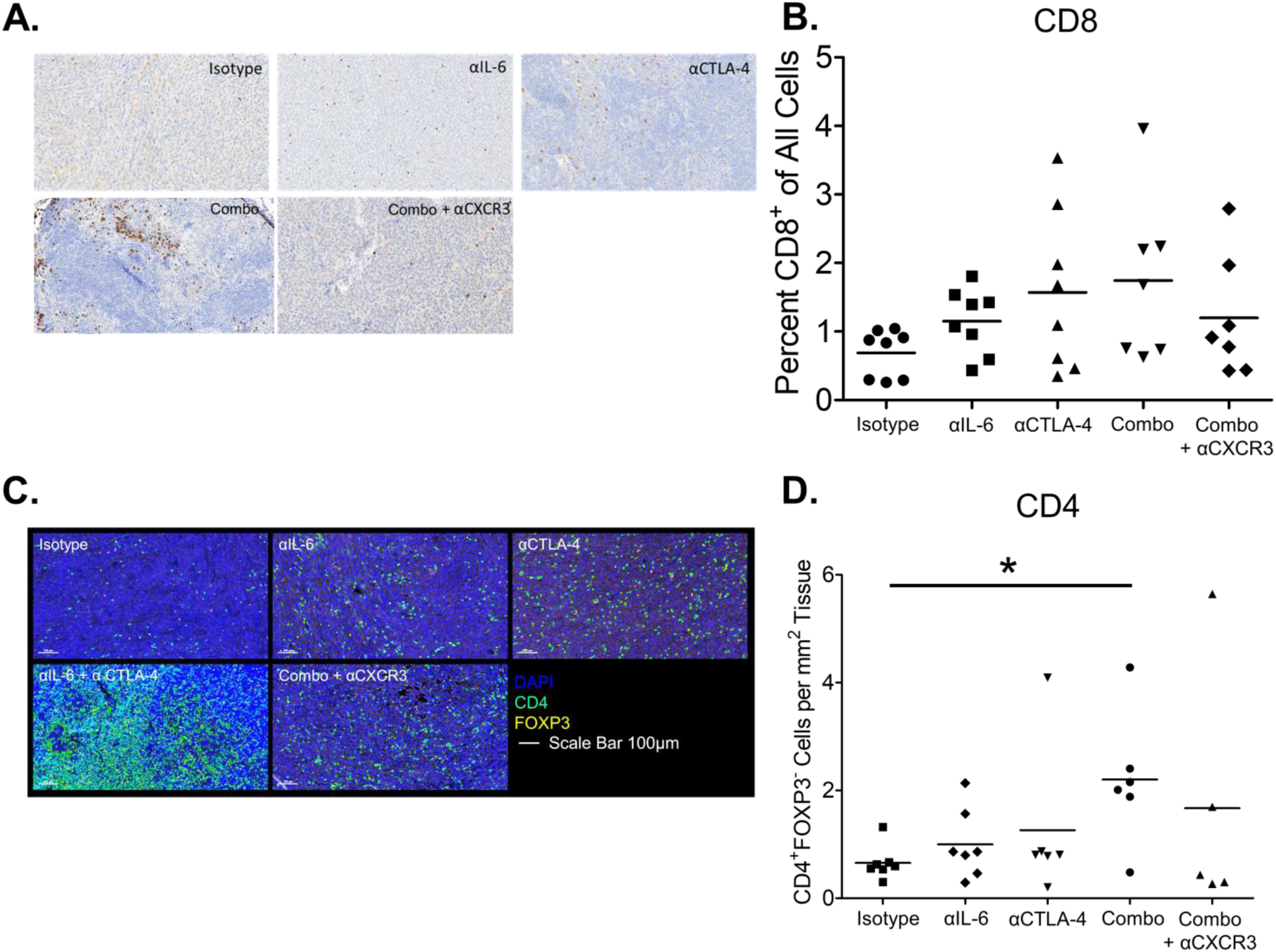
T cell infiltration of pancreatic tumors in altered in the presence of combined IL-6 and CTLA-4 blockade. **(A)** FFPE tissue slices of tumors from mice described in Figure 4 were stained for CD8 by IHC, quantified using Qupath and graphed as the percentage of cells expressing CD8. Results are displayed as mean for each treatment group. Symbols represent individual mice. **(B)** Representative images of IHC staining for CD8 in tumors from mice described in Figure 4A. **(C)** Slices from these tumors were also stained for CD4 and FOXP3 with DAPI counterstain followed by primary antibody-detection with Opal-conjugated antibodies. After scanning slides using a Perkin Elmer Vectra Polaris fluorescent slide scanner, the percentage of cells positive for CD4 but negative for FOXP3 were quantified with Qupath and graphed as mean for each treatment group. * indicates significant difference between isotype treated mice and mice receiving dual IL-6 and CTLA-4 blockade (P=0.0297). **(D)** Representative images are displayed with CD4 staining in green, FOXP3 staining in red and DAPI staining in blue. Scale bars are 100µm.

### Dual blockade therapy expands systemic TBET^+^ and GATA3^+^CD4^+^ T cell populations in a murine orthotopic PDAC model

Given the previous data demonstrating CD4^+^ T cells are required for the efficacy of this combination therapy, we next investigated the effects of this treatment on systemic T cell phenotypes. Flow cytometry staining and analysis (**Figure 6A**) recapitulated previous observations of Th1 immunity, features consistent with our splenocyte data in the subcutaneous model (**Figure 1D-E**). Robust expansion of CD3^+^CD4^+^TBET^+^ (Th1) T cells was observed in splenocytes from mice receiving combined IL-6 and CTLA-4 blockade (p=0.0015) or IL-6 blockade alone (p=0.0041) as compared to isotype treated mice (**Figure 6A**). We also observed a significantly higher frequency of splenic CD3^+^CD4^+^GATA3^+^ (Th2) T cells in mice treated with the combination compared to isotype treated mice (p=0.0003; **Figure 6B**). Of note, the CD3^+^CD4^+^GATA3^+^ T cells were more abundant in mice receiving the combination blockade, as compared to those treated with CTLA-4 blockade alone (p=.0012; **Figure 6B**). We also observed no difference in the frequency of T-regulatory cells, defined phenotypically as CD3^+^CD4^+^CD25^hi^FOXP3^+^, in splenocytes from mice receiving combined IL-6 and CTLA-4 blockade as compared to isotype control antibody treated mice (**Figure 6C**). Also, no difference in the numbers of CD3^+^CD4^+^RORγt^+^ cells was observed in mice receiving the combination compared to control mice (**Figure 6C**). However, single agent blockade of IL-6 resulted in a modest but significant (p=0.0088) increase in CD3^+^CD4^+^RORγt^+^ cells compared to isotype control antibody treated mice (**Figure 6C**). There were no differences in these systemic biomarkers between mice receiving only the combination or the combination together with CXCR3-targeted antibodies (**Figure 6A-D**).

**Figure 6.**
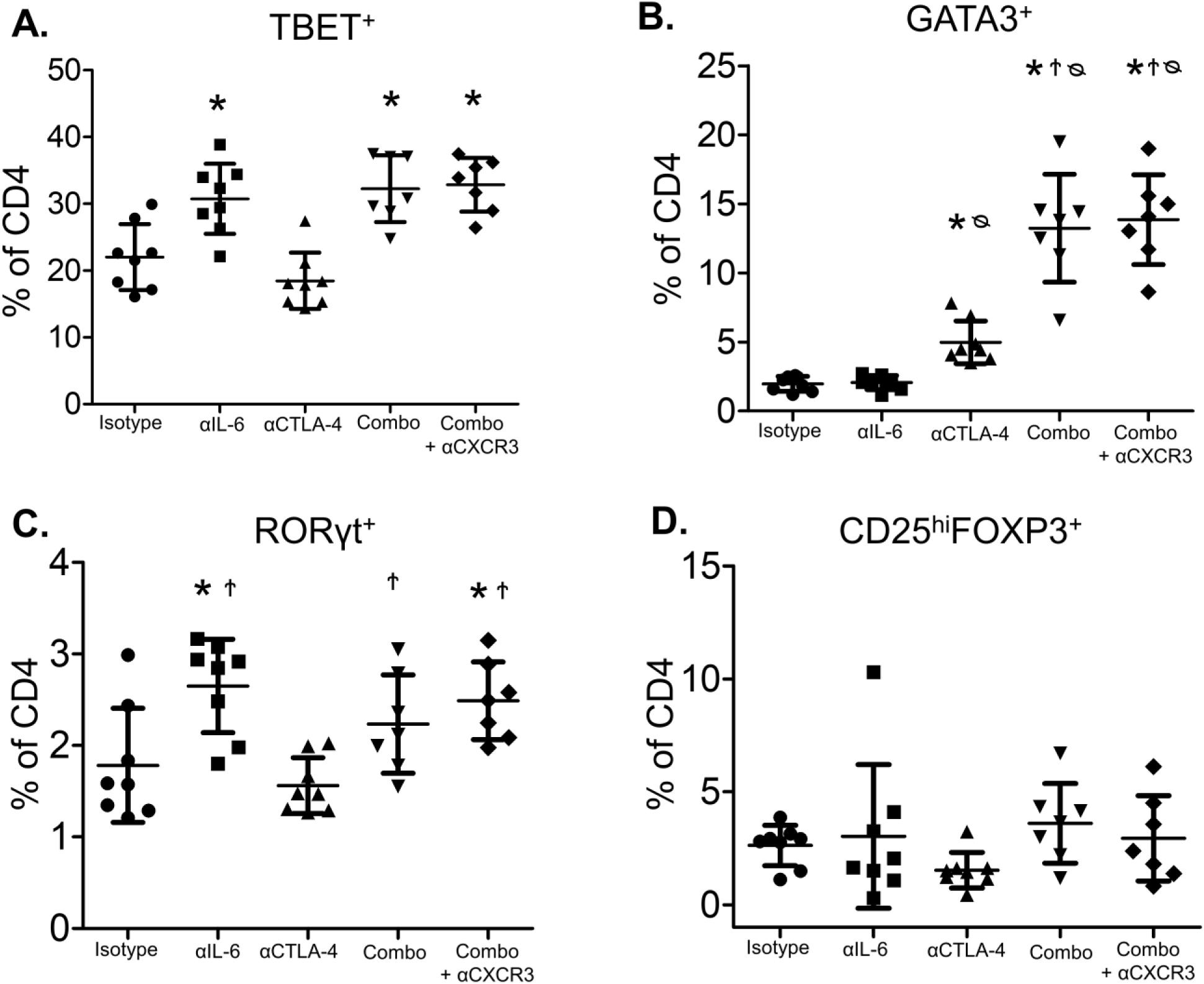
Combined blockade of IL-6 and CTLA-4 in mice bearing orthotopic pancreatic tumors results in systemic changes in CD4-helper T cells. Splenocytes were isolated from the mice receiving treatment as stated in Figure 4. Cells were stained for CD3, CD4, and CD8 surface markers with Ghost 780 viability stain to mark dead cells. After fixation and permeabilization, cells were stained for the transcription factors TBET, GATA3, FOXP3, and RORyt. The percentage of CD4^+^ T cells that were **(A)** TBET^+^ **(B)** GATA3^+^ **(C)** RORγt^+^ or **(D)** CD25^hi^FOXP3^+^ were graphed as mean ± SD with symbols indicating significance (p<0.05) compared to * isotype control mice, Ϯ αCTLA-4 treated mice, or ᴓ αIL-6 treated mice.

Additionally, we observed a higher percentage of CD4^+^ T cells positive for PD-1 in splenocytes from mice treated with IL-6 blockade alone or in combination with CTLA-4 blockade as compared to isotype control antibodies or single agent CTLA-4 blockade **(Supplemental Figure 4A).** Contrasting the data for CD4^+^ T cell subsets, few changes were observed in the composition of CD8^+^ T cells within the spleen of mice treated with single agent or combination therapy. However, mice receiving combined IL-6 and CTLA-4 blockade demonstrated significantly more CD8^+^ T cells expressing PD-1 as compared to mice receiving isotype control or anti-IL-6 antibodies (**Supplemental Figure 4B**). Immunologically, this therapeutic strategy dramatically impacted CD4^+^ T cells, driving increases in both Th1 and Th2 immunity, with more limited systemic changes in CD8^+^ T cells.

## Discussion

Antibodies that neutralize IL-6 have not been effective as single agent therapy in patients with pancreatic cancer (*27*). Moreover, blockade of CTLA-4 has limited efficacy as a single agent drug in this aggressive disease (*6, 28*). Here, we demonstrate that combined blockade of these targets elicits potent and long-lasting anti-tumor activity. Surprisingly, this dual therapy augmented treatment outcome in a CD4^+^ T cell-dependent manner. To our knowledge, this study represents the first attempt at this combination therapy approach in pancreatic cancer, as well as the first report to illuminate a mechanism that emphasizes the importance of CD4^+^ T cells in treatment outcome. We further define a mechanistic requirement for CD4^+^ T cells, along with the chemokine receptor CXCR3 in mediating T cell infiltration and resulting tumor regression. *In vitro*, dual blockade of IL-6 and CTLA-4 induced significant IFN-γ production by CD4^+^ T cells, while subsequent investigation revealed enhanced production of lymphocyte-specific chemokines by pancreatic tumor cells upon stimulation with IFN-γ. Together, our novel body of work highlight that dual IL-6 and CTLA-4 blockade alters chemokine production by pancreatic tumors, which likely fuels the trafficking of CD4^+^ T cells to the tumor (**Figure 7**).

**Figure 7.**
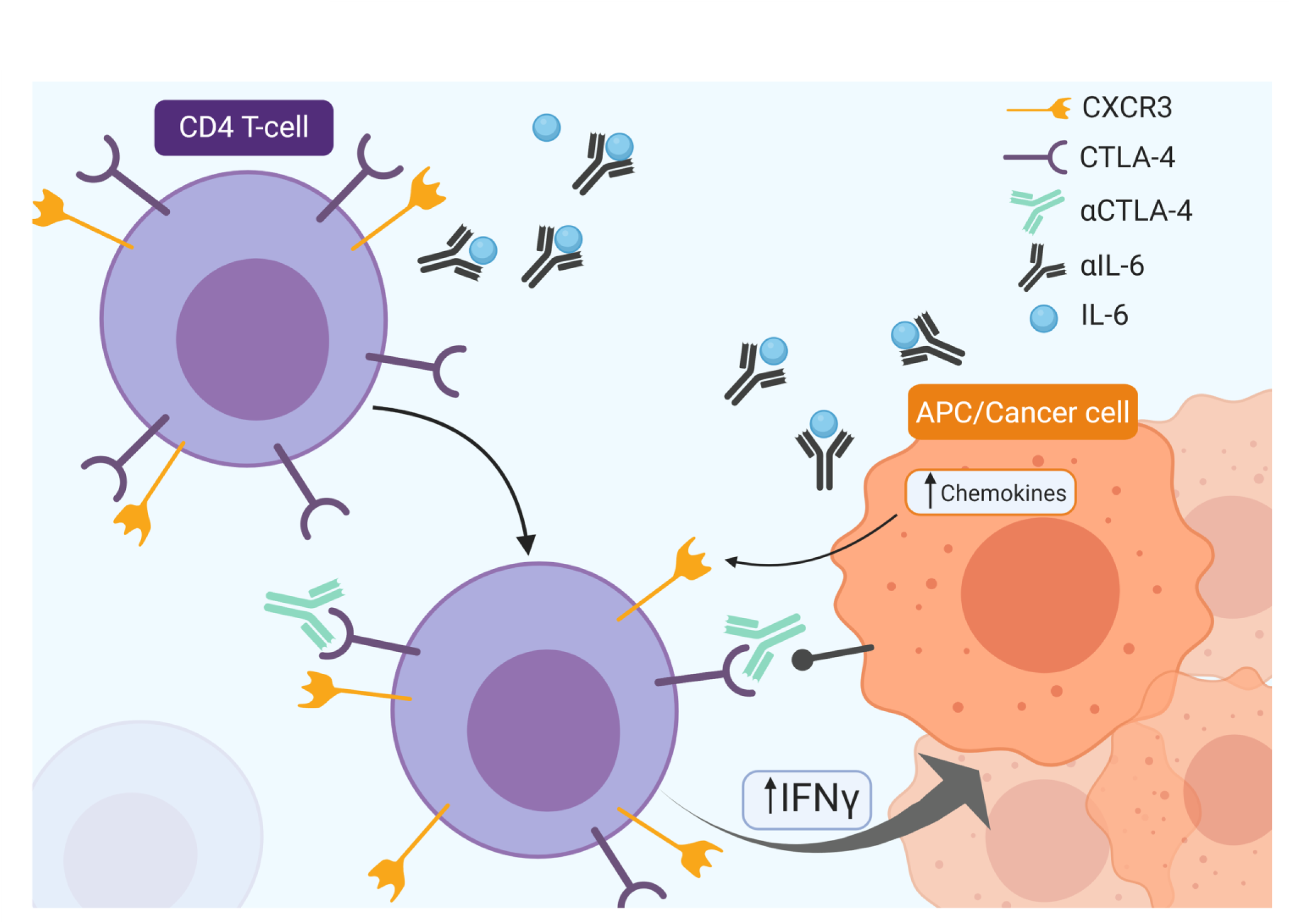
An illustrative representation of the proposed hypothesis by which IL-6 and CTLA-4 dual blockade therapy induces changes in T cell activity and subsequent production of chemokines to induce T cell migration into pancreatic tumors.

It is fascinating to consider how immune suppressive cytokines, produced by cancer associated fibroblasts and malignant cells, drive progression of this aggressive cancer and limit responses to immunotherapy. Initially, this idea was founded on our prior investigation of plasma from treatment-naive metastatic PDAC patients. This study revealed clues into systemic soluble factors and immune cells that might represent viable therapeutic targets: Namely, high serum IL-6 and circulating CTLA-4 expressing T cells were inversely correlated with overall survival (*13*). Complementing these data were earlier observations from our group and others that IL-6 is an abundant stromal-derived cytokine in pancreatic cancer, and may cooperate with other factors to promote the expansion of immunosuppressive myeloid cells (*16, 17, 29, 30*). The multi-faceted biologic effects of IL-6 on diverse cellular compartments (including tumor cells, fibroblasts, and immune cells) substantiates its influence as a key node in inflammation and cancer.

Targeting select cytokines, such as IL-6, has the unique potential to shift the overall immune response in a manner which alters Th1/Th2/Th17 cytokine balance that may be aberrantly regulated in advanced malignancy. In essence, this dysregulated cytokine balance can lead to direct inhibition of cytotoxic effector cell function; 2) expansion of immune suppressive lymphoid and myeloid cell populations; 3) reciprocal activation of pro-oncogenic and pro-metastatic pathways in the tumor cells (*16, 20-22, 31-34*). In the setting of PDAC, the stroma contributes significantly to the production of tumor-promoting cytokines and chemokines. Our group has found the cytokine IL-6 to be a particularly important mediator of interactions between the stroma, tumor and immune system in PDAC (*13, 16, 17*). We view the strategy of targeting IL-6 as a novel approach with promise to unlock enhanced efficacy of ICI and other targeted therapies in traditionally resistant diseases such as PDAC.

The efficacy of concurrently targeting IL-6 and T cell intrinsic immune checkpoints was first established by our group in murine PDAC models with PD-1/PD-L1 blockade (*17*). Strikingly, this effect has since been reproduced in multiple murine solid tumor models including glioblastoma, colorectal cancer and melanoma (*35–37*). The data from this current report is the first to indicate that IL-6 blockade also augments the efficacy of antibodies targeting CTLA-4 in PDAC. These results are particularly important in lending flexibility to clinical translation whereby multiple immune checkpoint antibodies may have efficacy via non-overlapping mechanisms of action. Results from this study indicate a unique mechanism of action when compared to combined IL-6 and PD-1/PD-L1 blockade, with several evident differences. Most notable is the reliance of IL-6 and PD-1/PD-L1 blockade on CD8^+^ T cells while IL-6 and CTLA-4 blockade is clearly CD4^+^ T cell dependent. Additionally, blockade of IL-6 and PD-1/PD-L1 reduced stromal content, while CTLA-4 and IL-6 revealed no such changes as determined by histological staining of murine tumors for the activated fibroblast marker alpha-smooth muscle actin (αSMA) (*17*). This possible discrepancy may be due to differential effects on various subsets of cancer-associated fibroblasts that could be affected by therapy, one of which produces high levels of IL-6 (*38, 39*). These results highlighted a unique dependence on CD4^+^ T cells for efficacy of this combined therapy.

Previous studies investigating T cell responses to pancreatic cancer have proposed differing results with respect to the effect of CTLA-4 on CD4^+^ or CD8^+^ T cells in pancreatic cancer (*19, 40*). Here depletion of CD4^+^ or CD8^+^ T cells indicates a heavy dependence on CD4^+^ T cells to mediate the anti-tumor effects of combined IL-6 and CTLA-4 blockade, with CD8 depletion only moderately restoring tumor growth. CD4^+^ T-helper support of CD8 responses in the pancreatic tumor microenvironment may provide heterogeneous T cell responses that effectively kill neoplastic cells and mediate tumor regression. Recent evidence from the Schreiber group describes the need for CD4^+^ and CD8^+^ T cell activation for effective anti-tumor responses (*41*). In particular, the response of the host to immune-based therapies requires CD4 responses to mediate effective T cell-based rejection of tumors. Here, our studies reinforce the emerging importance of CD4^+^ T cells in mediating efficacy of immune therapies. As immunotherapy combination approaches move into the clinical setting, it will be important to grasp how these therapies impact the various T cell subsets present in the tumor microenvironment.

We hypothesized that the efficacy of IL-6 and CTLA-4 blockade may be mediated in part by altering the balance of chemokines and subsequent trafficking of lymphocyte populations into the tumor microenvironment. IL-6 has previously been shown to regulate the percentage of lymphocytes expressing receptors such as CXCR3 that are associated with CD8 and Th1 recruitment to sites of inflammation (*42*). Additionally, studies of single agent CTLA-4 blockade in murine models of pancreatic cancer have reported effects of single agent CTLA-4 blockade on CD4^+^ T cell infiltration without generating the dramatic tumor regression observed here (*19*). The tumor protective effects of CXCR3 blockade observed in this report suggests that infiltrating Th1-helper cells and CD8^+^ T cells may contribute to tumor regression in the context of this combination therapy. Indeed, recent reports support the importance of the CXCR3 chemokine axis for mediating tumor responses to ICI (*43*), supporting our observations here.

Conversely, the expansion of cells expressing the Th2 transcription factor GATA3 was somewhat unexpected. The systemic changes in Th2-helper cells, defined by GATA3 expression, and the influence of IL-6 on tumor-derived chemokines suggests an effect on CD4^+^ T cells other than those of the Th1-helper subset. A review of the literature supports this response, as others have demonstrated CTLA-4 blockade to induce the expansion of Th2-helper CD4 cells in the context of cancer (*44, 45*). We propose that while blocking IL-6 in our murine models of pancreatic cancer drives Th1-helper responses, anti-IL-6 antibodies may also alleviate mechanisms preventing the expansion of other T cell subsets.

While this combination therapy elicits potent and efficacious anti-tumor activity, clinical application of ICI has potential to induce toxicity for some patients. Attempts to ameliorate the autoimmune toxicities of ICI including anti-CTLA-4 and anti-PD-1/PD-L1 targeted antibodies have revealed the IL-6R blocking antibody tocilizumab is effective in patients refractory to steroids (*12, 46*). A recent report demonstrated the IL-6R-blocking antibody tocilizumab alongside pembrolizumab in the treatment of advanced melanoma not only prevented exacerbation of Crohn’s disease, but also allowed for a durable anti-tumor immune response (*12*). Similarly, a recent case report showed IL-6 blockade with tocilizumab in a patient with stage IV pulmonary adenocarcinoma completely resolved immune related toxicities to nivolumab including oropharyngeal mucositis and esophagitis with severe esophageal stenosis (*47*). Finally, the use of IL-6R-blocking antibodies with chimeric antigen receptor T cell (CAR T) therapy has produced encouraging results while actually enhancing patient safety (*46*). This emerging use of IL-6/IL-6R blockade to limit ICI-associated toxicities has actually led to clinical trials exploring the use of IL-6 blockade specifically for improved safety of these therapeutic options (NCT03601611).

Despite these encouraging results, efforts to apply IL-6 or IL-6R blockade prospectively with therapeutic intent in clinical trials have lagged behind the pre-clinical data, possibly due to resistance to repurpose these drugs from the autoimmunity space into the oncology setting. To date, only a single clinical trial from our group (NCT04191421) is exploring combining IL-6 blockade therapy with ICI for the treatment of any oncology indication (pancreatic cancer). Continued clinical experience with IL-6 and ICI combinations across solid tumors will inform the field regarding both their efficacy and ability to limit autoimmune sequelae, moving the field forward.

## Materials and Methods

### Cell lines and antibodies

Murine MT5 (Kras^LSL-G12D^, Trp53^LSL-R270H^, Pdx1-cre) pancreatic cells were a gift from David Tuveson (Cold Spring Harbor Laboratory, Cold Spring Harbor, NY) and cultured in RPMI-1640 (Gibco) with 10% FBS, 10 mM L-glutamine, and antibiotics (GiminiBio). Murine KPC-luc (Kras^LSL-R270H^, p53^-/-^, Pdx1-cre) cells expressing an enhanced firefly luciferase construct (a gift from Dr. Craig Logsdon, MD Anderson Cancer Center) were cultured in DMEM (Gibco) with 10% FBS, 10nM L-glutamine and antibiotics. Murine antibodies to IL-6 (Clone MP5-20F3), CTLA-4 (Clone 9D9), CXCR3 (Clone CXCR3-173) or isotype controls (Clones LTF-2 for subcutaneous or HRPN for orthotopic studies, MCP-11 and polyclonal Armeniam hamster IgG, respectively) were purchased from BioXcell (West Lebanon, NH) for *in vivo* studies. Additionally, anti-mouse CD8a (Clone 2.43) and anti-mouse CD4 (Clone GK1.5) depleting antibodies were purchased from BioXcell.

### In vivo murine efficacy studies

All animal studies were conducted under an approved institutional animal care and use committee (IACUC) at The Ohio State University or Emory University. For subcutaneous *in vivo* efficacy studies, 1×10^6^ MT5 tumor cells were injected subcutaneously in the flank of female C57BL/6 mice. Once tumors reached 50 – 100 mm^3^ (typically 7-10 days), antibody treatment was initiated. Subcutaneous studies were ended once tumors in mice receiving control antibodies reached a volume that meet IACUC-mandated early removal criteria. For orthotopic *in vivo* efficacy studies, 2×10^5^ KPC-luc tumor cells were orthotopically injected into the pancreas of 6-8 week old female C57BL/6 mice. Tumors were permitted to grow for 7 days prior to randomization into treatment group and presence of pancreatic tumors were confirmed by bioluminescent imaging as previously described (*17*). Animals were treated 3 times each week with 200µg/mouse of isotype, anti-IL-6 or anti-CTLA-4 antibodies (BioXCell). For CXCR3 blockade studies, anti-CXCR3 antibody or appropriate isotype control antibody were administered concurrently with therapeutic antibody according to the same schedule at 200µg/mouse until completion of the study as described previously (*48*). For *in vivo* studies incorporating antibody-based depletion of CD4^+^ or CD8^+^ T cells, animals were treated with 200µg/mouse of antibodies for CD4 or CD8 depletion on a schedule as previously described (*49*), with the alteration that animals were treated every 3 days, rather than every 4 days, after the first week.

### Antibodies for flow cytometry and immunohistochemical staining

A list of all antibodies, with clone names, used for flow cytometry and immunohistochemistry can be found in supplementary table 1.

### Flow cytometry

At the completion of subcutaneous *in vivo* efficacy studies, tissues were harvested for subsequent immunophenotypic analyses of splenocytes and single cell suspensions from tumors were assessed by flow cytometry as previously described (*17*). Briefly, antibodies for MDSC were CD11b, Ly6G, and Ly6C. Tregs were defined by staining for CD4, CD25 and FOXP3 and Th17 cells were defined by staining for CD4 and RORγt. For analysis of T cell activation markers, cells were stained with Ab specific for CD4. CD8, CD62L, and CD44. To determine Th1 or Th2 phenotypes, cells were stained using antibodies to CXCR3, CCR4, and CCR6. Cells were incubated on ice for 30 minutes, washed, and fixed in PBS containing 1% formalin for flow cytometric analysis on a LSRII flow cytometer (BD Biosciences). For FOXP3 and RORγt staining, cells stained for surface markers were fixed and permeabilized for staining of transcription factors using the eBiosceince Foxp3/Transcription Factor Staining Buffer Set according to manufacturer’s suggested protocol. Flow cytometry analysis of splenocytes from orthotopic studies identified Th1-helper cells as CD3^+^CD4^+^CD8^-^TBET^+,^ Th2-helper cells as CD3^+^CD4^+^CD8^-^GATA3^+^, Th17-helper cells as CD3^+^CD4^+^CD8^-^RORγt^+^ and Tregs as CD3^+^CD4^+^CD8^-^CD25^high^FOXP3^+^. These subsets were further analyzed for surface expression of PD-1 and ICOS. Within the population of CD3^+^CD8^+^CD4^-^ cells, we identified stem-like cells as PD-1^+^, TCF1/7^+^ and TIM3^-^ and effectors as PD-1^+^, TCF1/7^-^ and TIM3^+^. Additionally, the expression of CTLA-4, CD69, CD39 and CD44 were analyzed. Cells were also stained with Ghost 780 viability dye to detect live cells. After surface staining, cells were fixed and permeabilized for staining of transcription factors using the eBiosceince Foxp3/Transcription Factor Staining Buffer Set according to the manufacturer’s suggested protocol. Samples were then run on a Cytek Aurora flow cytometer (Cytek).

### Immunohistochemical analysis

Formalin fixed paraffin-embedded (FFPE) tumors from subcutaneous *in vivo* experiments in mice were subjected to IHC analysis following staining with Ab against CD3 (Catalog A0452; Dako). For quantification, 40x magnification images of tumors (10 images per mouse tumor) were captured using PerkinElmer’s Vectra multispectral slide analysis system. inForm software tools were used to quantify CD3-positive cells (Fast Red chromogen) within each image. Additional tissue slices were stained with antibodies against CD8, and αSMA and scanned with an Olympus Nanozoomer whole slide scanner and analyzed using Qupath (CD8) or FIJI (NIH) in the case of αSMA. Livers and tumors from mice bearing orthotopic tumors were also FFPE. Dual stains for DAPI (Perkin Elmer) with CD4 and FOXP3 were performed using a Roche autostainer. Opal 520 conjugated secondary (Perkin Elmer) was used to detect CD4 and Opal 630 (Perkin Elmer) conjugated secondary was used to detect FOXP3. Multiplex stained slides were imaged using the Vectra Multispectral Imaging System version 2 (Perkin Elmer). Filter cubes used for multispectral imaging were DAPI (440–680 nm), FITC (520 nm-680 nm), Cy3 (570–690 nm), Texas Red (580– 700 nm) and Cy5 (670–720 nm). Multispectral images were then analyzed with Qupath (*50*).

### TRP-1 transgenic CD4^+^ T cell activation

CD4^+^ TRP-1 transgenic T cells (*51*) were activated with TRP-1106-130 peptide (SGHNCGTCRPGWRGAACNQKILTVR) loaded at a 1μM concentration onto irradiated B6 splenocytes (10Gy) at a 2:1 TRP-1:feeder cell ratio. TRP-1 cells were cultured in the presence of monoclonal antibodies targeting CTLA-4 (10μg/mL, clone 9D9), IL-6 (10μg/mL, clone MP5-20F3) or Isotype controls (10μg/mL, IgG2b or HRPN) as indicated with IL-2 at 100IU/mL. Cells were assessed three days after activation for cytokine production post PMA/Ionomycin stimulation. Briefly, cells were activated in PMA (30nM) and Ionomycin (20nM) (Sigma) with Monensin (2μM) and Brefeldin A (5μg/mL) (Biolegend) for 4 hours, followed by fixation and permeabilization for cytokine staining according to manufacturer protocol (BioLegend)

### In vitro evaluation of chemokine production

KPC-luc cells were plated at 2×10^5^ cells per well in 6 well plates. Media was then supplemented with 10ng/ml IL-6 (peprotech), 1μg/ml IFNy (peprotech), both cytokines combined, or vehicle control for 24 hours. Supernatant was then collected and spun at 1000 *x g* before being transferred to a new tube to ensure no cellular contamination. Supernatants were analyzed for relative changes in chemokine levels using the Proteome Profiler Mouse Chemokine Array Kit (ARY020, R&D Systems). Results from this array were confirmed using DuoSet ELISA kits from R&D Systems for CXCL10, CXCL9, CCL2, and CCL5. MT5 cells were also plated at 2×10^5^ cells per well and supernatants collected in the same fashion as above. The same ELISA kits for CXCL10, CXCL9, CCL2 and CCL5 were used to evaluate conditioned supernatants from these experiments as well.

### Statistical Analysis

Data from subcutaneous studies obtained by flow cytometry and IHC as well as tumor volumes were log-transformed prior to analysis to meet model assumptions of normality and homoscedasticity. Tumor volume was modeled over time using mixed-effects regression with fixed effects for group, time and the interaction of the two. Random intercepts and slopes by mouse were included with an unstructured covariance matrix for the random effects. Other outcomes were compared using ANOVA. P-values of less than 0.05 were considered significant.

For *in vivo* orthotopic studies, descriptive statistics for each variable were reported. For numeric covariates, the mean and standard deviation were calculated and presented. One-way ANOVA was performed for IHC data and splenocyte data with univariate analysis. Least significant difference method (LSD) was used for pairwise multiple comparisons. Natural log transformation of bioluminescent imaging data was performed to achieve approximately normal distribution of data. For log transformed data, linear mixed models were used to test whether there was any significant change over time of each outcome and to detect whether there is any significant difference of each outcome among different treatments. The significance level was set at 0.05. For *in vitro* data, natural log transformation was performed to normally distribute data and we then performed one-way ANOVA and LSD to detect whether means significantly differed among the treatment groups. All analyses were conducted in SAS v9.4 (SAS Institute, Cary, NC).

## Acknowledgements

We would like to acknowledge the cores at Winship Cancer Institute and Emory University that made this research possible including the Pediatric/Winship Flow Cytometry Core, the Winship Cancer Tissue and Pathology Shared Resource, the Winship Biostatistics and Bioinformatics Core, the Winship Cancer Animal Models Shared Resource and the Winship Integrated Cellular Imaging Core and the Emory Integrated Genomics Core (EIGC), under NIH/NCI award number P30CA138292. The content is solely the responsibility of the authors and does not necessarily represent the official views of the National Institutes of Health.

