## Supplementary material for "Dual blockade of IL-6 and CTLA-4 regresses pancreatic tumors in a CD4^+^ T cell-dependent manner": Suplemental Figures

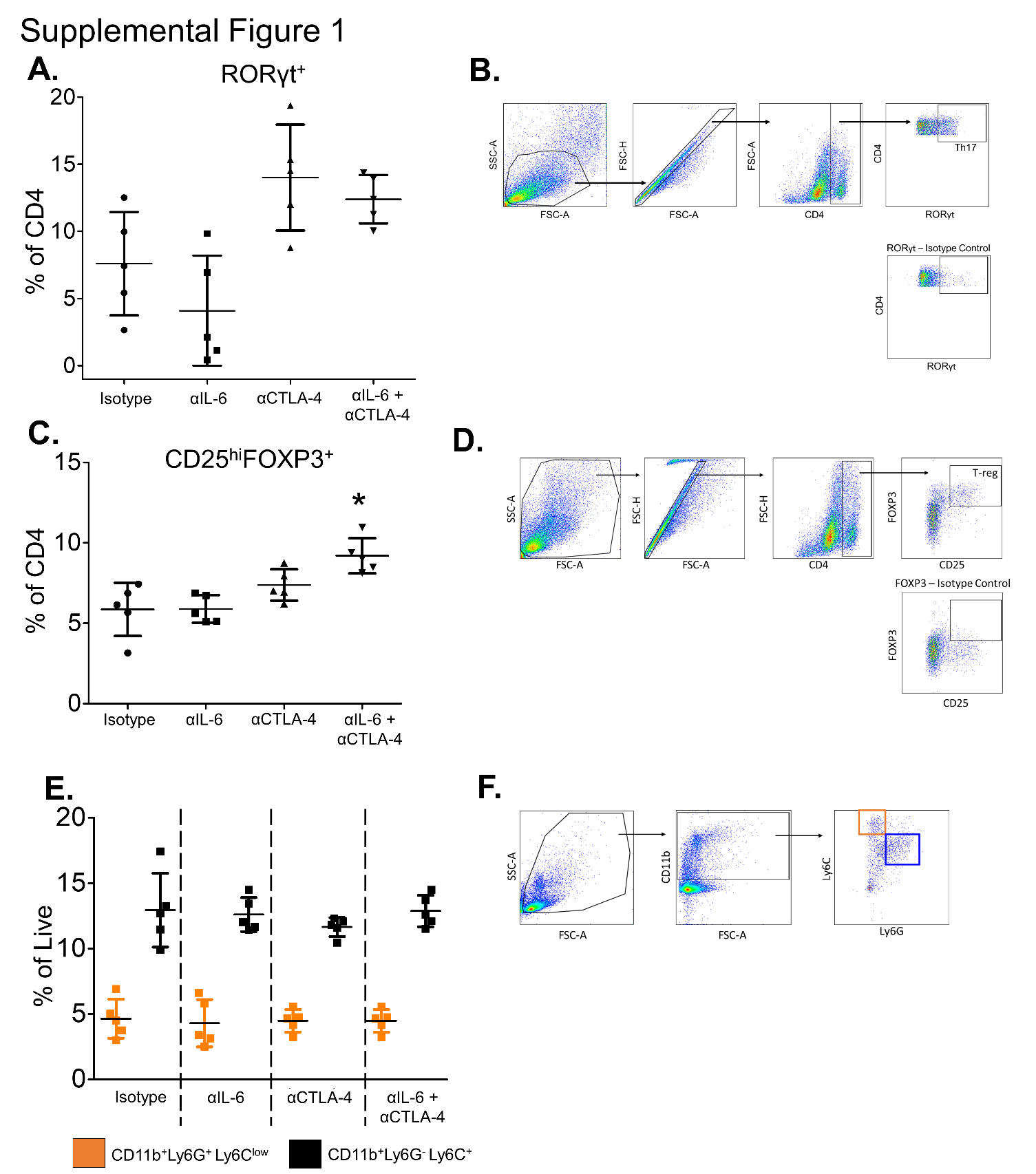

**Supplemental Figure 1.** Splenocytes were isolated from the spleens of mice from the subcutaneous therapeutic study described in **Figure 1. (A)** Splenocytes were stained with antibodies to CD4, CD25 and FOXP3 and analyzed by flow cytometry. The percentage of CD4^+^ cells that were CD25^hi^FOXP3^+^ were graphed as mean ± SD. * indicates significance (P<0.05) compared to isotype control treated animals. **(B)** Gating strategy for the identification of Tregs (CD4^+^CD25^hi^FOXP3^+^) **(C)** Splenocytes were stained with antibodies to CD4 and RORγt and analyzed by flow cytometry. The percentage of CD4^+^ cells that were RORγt^+^ were graphed as mean ± SD. **(D)** Gating strategy for the identification of Th17 cells (CD4^+^RORγt^+^) **(E)** Splenocytes were also stained with antibodies to CD11b, Ly6C and Ly6G and analyzed by flow cytometry. The percentage of CD4^+^ cells that were RORγt^+^ were graphed as mean ± SD. **(F)** Gating strategy for the identification of monocytic-MDSCs (CD11b^+^Ly6G^-^Ly6C^+^) and polymorphonuclear-MDSCs (CD11b^+^Ly6G^+^Ly6C^lo^).

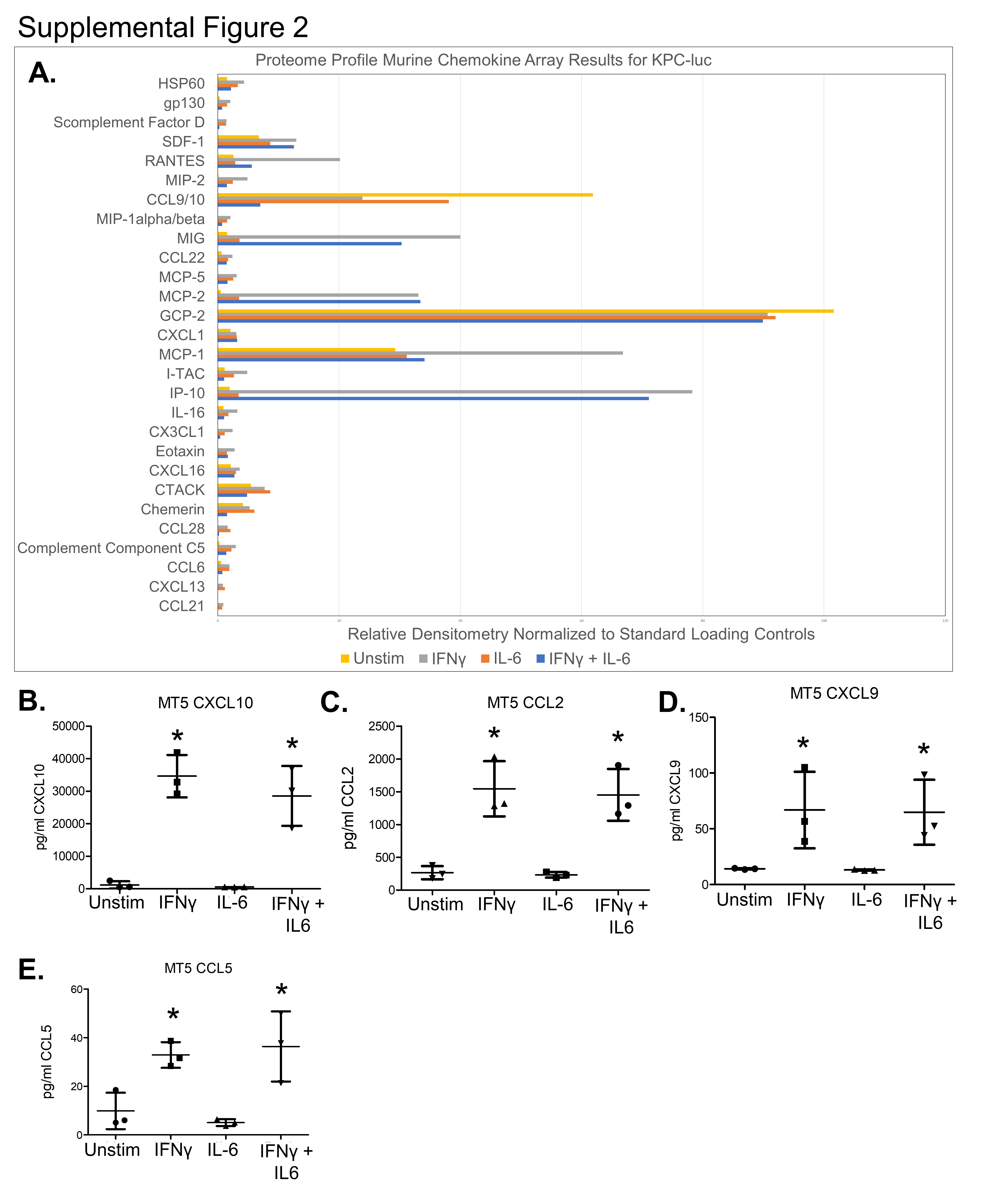

**Supplemental Figure 2. (A)** Relative densitometry for each chemokine produced by KPC-luc cells unstimulated (yellow), or treated with 1μg/ml IFNγ (grey), 10ng/ml IL-6 (Orange), or the combination of IFNγ and IL-6 (Blue). ELISA quantification of **(B)** CXCL10 **(C)** CCL2 **(D)** CXCL9 and **(E)** CCL5 production by MT5 pancreatic cancer cells unstimulated or treated with 1μg/ml IFNγ, 10ng/ml IL-6, or the combination of both IFNγ and IL-6. Data is graphed as mean of 3 replicates ± SD. * indicates significance (p<0.05) compared to unstimulated cells or cells treated with IL-6.

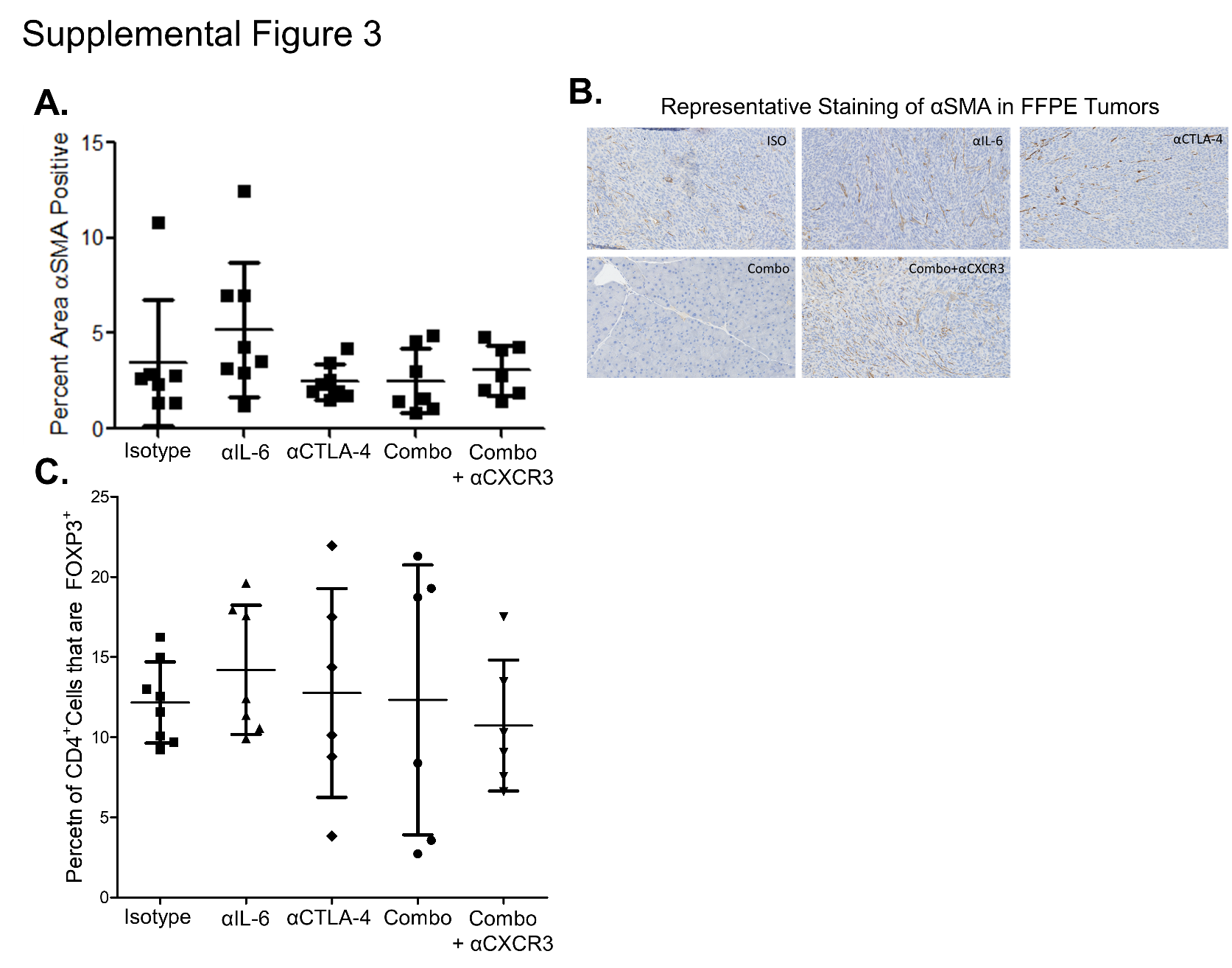

**Supplemental Figure 3.** Tissue slices of FFPE tumors from mice in the orthotopic study outlined in Figure 4 were stained for alpha-smooth muscle actin (αSMA) or CD4, FOXP3 and DAPI. **(A)** Each tissue slice stained for αSMA was sampled based on total tissue area and the resulting 20x images were analyzed using FIJI to determine the percent area positive for αSMA. The mean ± SD was then graphed for each treatment group. **(B)** Representative 20x images of αSMA staining of tumors from each treatment group. **(C)** Immunofluorescent whole slide scans were collected on the Vectra Polaris Slide Scanner and analyzed using the Qupath to determine the percentage of CD4^+^ cells expressing FOXP3. The mean ± SD for each group was graphed.

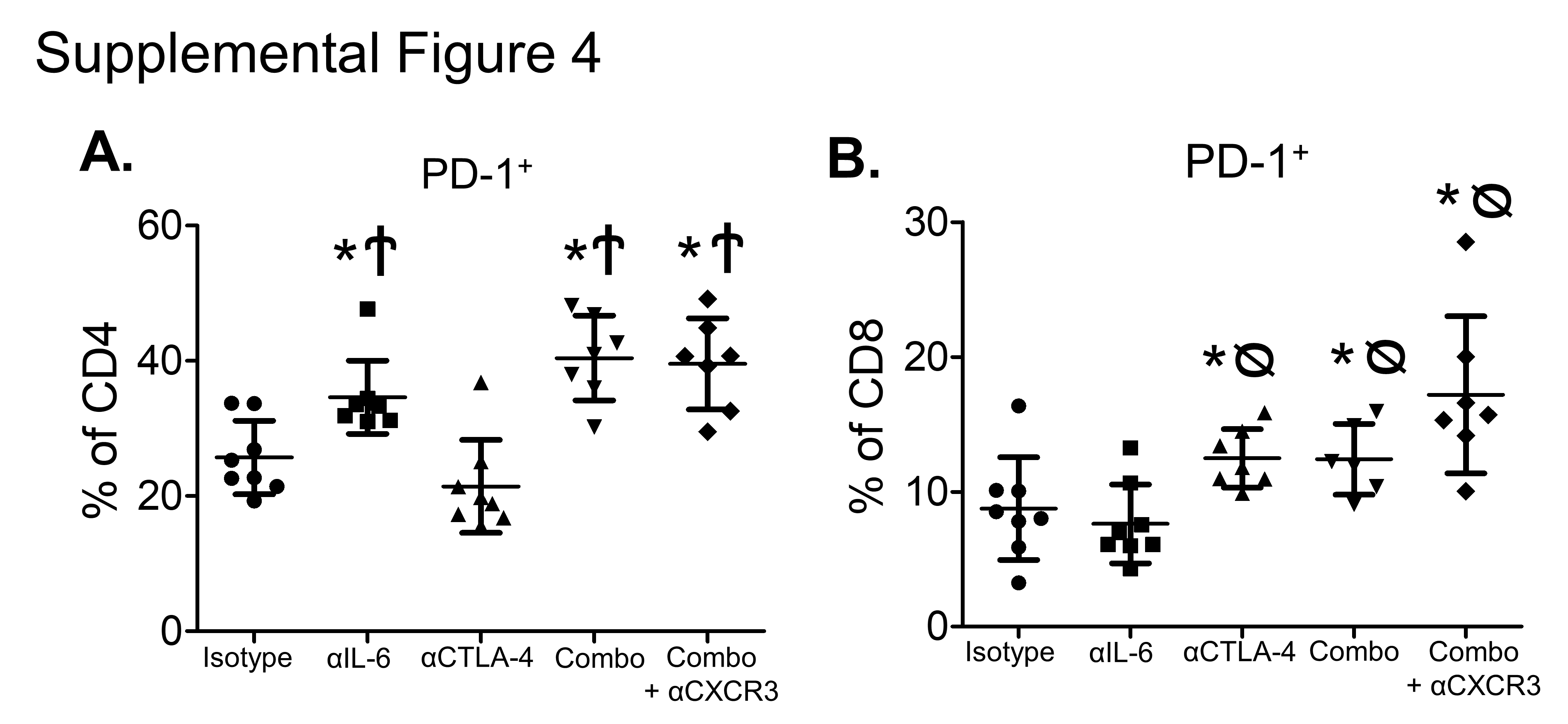

**Supplemental Figure 4.** Splenocytes isolated from mice treated in the orthotopic study described in Figure 4 were stained and the percentage of **(A)** CD8^+^ T cells expressing PD-1 or **(B)** CD4^+^ T cells expressing PD-1 were determined by flow cytometry analysis. Graphs show mean ± SD for each treatment group. * indicates significance (p<0.05) to isotype control treated mice, ᴓ indicates significance (p<0.05) to mice treated with IL-6 antibody, and Ϯ indicates significance (p<0.05) to CTLA-4 antibody treated mice.

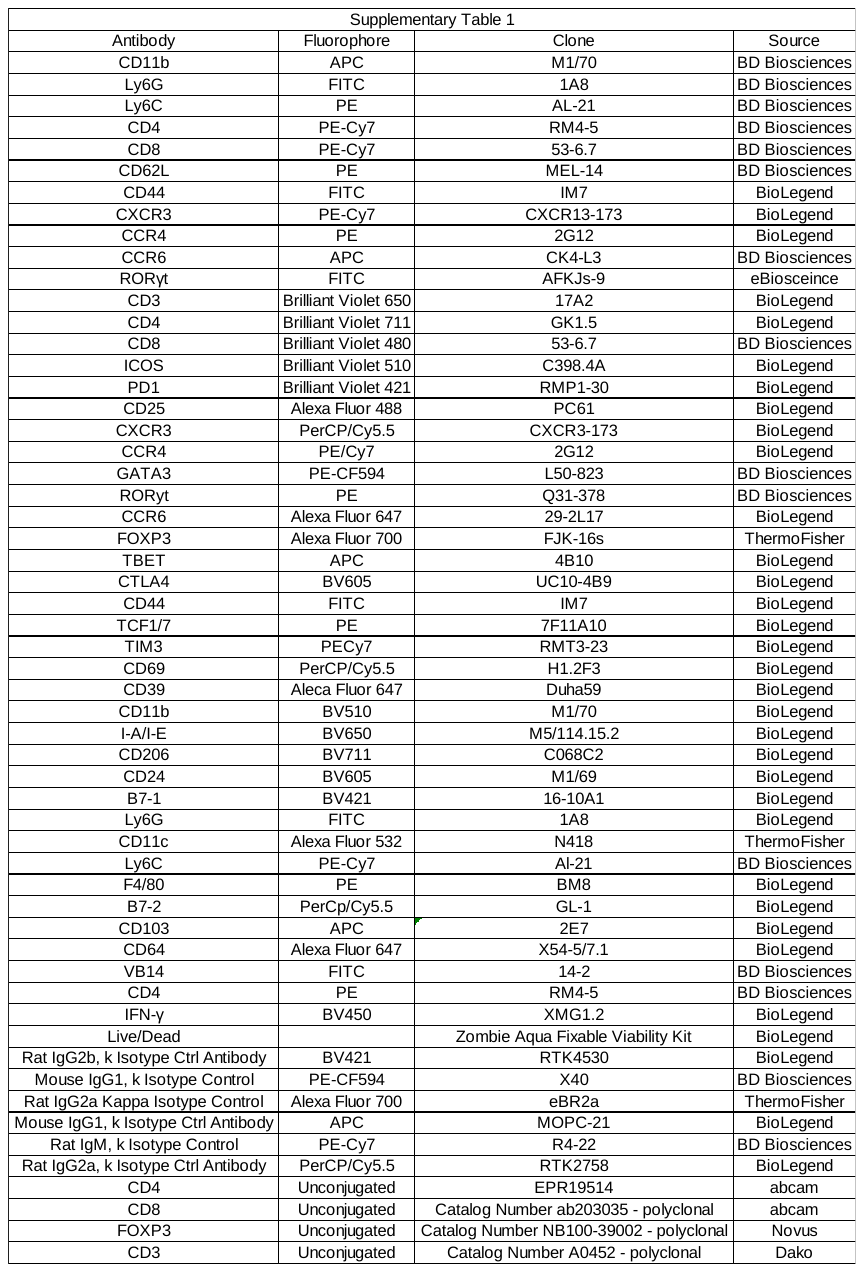

**Supplemental Table 1**. A list of all antibodies used for flow cytometry and imaging.
